# Single-cell RNA sequencing of iPSC-derived brain organoids reveals *Treponema pallidum* infection inhibiting neurodevelopment

**DOI:** 10.1101/2024.01.23.576898

**Authors:** Qiu-Yan Xu, Yong-Jing Wang, Yun He, Xin-Qi Zheng, Man-Li Tong, Yu Lin, Tian-Ci Yang

**Affiliations:** Centre of Clinical Laboratory, Zhongshan Hospital of Xiamen University, School of Medicine, Xiamen University, Xiamen, 361004, China; Guangyuan Hospital of Traditional Chinese Medicine, Guangyuan, 628000, China; Department of Medical Laboratory, The Second Affiliated Hospital of Xiamen Medical College, Xiamen, 361021, China; Institute of Infectious Disease, School of Medicine, Xiamen University, Xiamen, 361004, China

**Keywords:** Congenital syphilis, *Treponema pallidum*, Brain organoids, Single-cell RNA sequencing, Neurodevelopment

## Abstract

Congenital syphilis is a vertically transmitted bacterial infection caused by *Treponema pallidum*, often causing multidomain neurodevelopmental disabilities. However, little is known about the pathogenesis of this disease. Brain organoids platform derived from the induced pluripotent stem cell (iPSC) is exposed to *T. pallidum* infection for modelling congenital neurodevelopmental impairment. Single-cell RNA sequencing is used for identifying the subpopulations of differentially expressed genes and cellular heterogeneity and reconstructing differentiation trajectories following *T. pallidum* infection. The results reveal that *T. pallidum* infection influences the formation of neural rosette structures, reduces the cell number of the neural progenitor cell subcluster 1B (subNPC1B) and hindbrain neurons, and affects the neurodevelopment of the brain organoid. Moreover, it is speculated that *T. pallidum* inhibits the hindbrain neuron cell number through the suppression of subNPC1B subgroup in the organoids and inhibits transcription factor 3 activity in the subNPC1B-hindbrain neuronal axis. This is the first report on the inhibited effects of *T. pallidum* on the neurodevelopment of the iPSC-derived brain organoid model. *T. pallidum* could inhibit the differentiation of subNPC1B in brain organoids, thereby reducing the differentiation from subNPC1B to hindbrain neurons, and ultimately affecting the development and maturation of hindbrain neurons.

## Introduction

Congenital infections, which refer to maternal infections with vertical transmission to the fetus, pose a significant global public health issue and are associated with adverse pregnancy outcomes, neonatal diseases, long-term neurodevelopmental consequences, and increased healthcare expenditures (Grosse et al., 2021). Notably, an expanding roster of congenital infections, such as cytomegalovirus, Zika virus, syphilis, rubella, lymphocytic choriomeningitis virus, and toxoplasmosis, have been implicated in neurodevelopmental impairments (Fortin and Mulkey, 2023). With the increasing number of syphilis cases in recent years in many countries, there has been a growing focus on the prenatal exposure of the foetus to maternal *Treponema pallidum* and its consequential effects on the developing brain and subsequent neurodevelopmental outcomes (David et al., 2022; Medoro and Sánchez, 2021). Tonni et al. upon utilising a transabdominal neuroscan of a pregnant woman with primary syphilis observed hyperechogenicity involving both choroid plexuses of the lateral ventricles and the falx cerebri (Tonni et al., 2022). Moreover, the foetal brain contour exhibited a smooth appearance devoid of discernible gyri and sulci, indicating a significant influence of *T. pallidum* on fetal brain development and subsequent neurodevelopmental outcomes (Tonni et al., 2022). However, the precise mechanisms underlying the impact of *T. pallidum* on fetal neurodevelopment remain constrained.

Self-renewing and pluripotency properties of human pluripotent stem cells have greatly facilitated the understanding of the human nervous system development and pathogenesis of various neurological diseases (Mertens et al., 2016; Ramani et al., 2020). Human brain organoids derived from induced pluripotent stem cells (iPSCs), which more closely resemble the developing human brain than previous approaches (Lancaster et al., 2013), provide unprecedented opportunities for studying viral infections and their impact on human neurodevelopment (Kim et al., 2019; Krenn et al., 2021). The development of organoids requires robust strategies to assess the cellular composition of organoids and their complex interactions and neuronal networks. Single-cell RNA sequencing (scRNA-seq), a high-resolution approach that fits the above requirement, can deconstruct organoid cell composition, help study the molecular underpinnings of cellular heterogeneity, and reconstruct differentiation trajectories in these complex developing tissues (Kanton et al., 2019; Velasco et al., 2019). Several laboratories using scRNA-seq for the direct comparison of cultured and primary cortical structures *in vitro* and *in vivo*, respectively, demonstrated great similarity in the gene expression profiles (Camp et al., 2015; Pollen et al., 2019). However, understanding and interpreting gene expression and regulatory patterns from single-cell genomic data on organoids in the field of syphilis remains a major challenge. Here, we constructed the iPSC-derived brain organoids and infected them with *T. pallidum*, and evaluated the neurodevelopment of the brain organoids based on the scRNA-seq to model and analyse the pathogenesis of neurodevelopmental disorders of congenital syphilis. This is the first study to evaluate the effects of *T. pallidum* on the neural development of an iPSC-derived brain organoid model, thereby laying the foundation for further research elucidating the pathogenesis of congenital syphilis.

## Results

### *T. pallidum* infection inhibited brain organoid neurodevelopment

To study the neurodevelopment of *T. pallidum*-infected brain organoids instead of patient-derived brain tissue, we first generated iPSC-derived brain organoids for nearly 2 months as previously described (Madhavan et al., 2018; Setoh et al., 2019) (Figure 1A, Supplement file-figure supplement 1). Immature brain organoids on day 25, which coincides with the gradual differentiation and maturation of brain organoid, were utilised for *T. pallidum* infection. Brain organoid growth was tracked over 30 days post-infection to monitor organoid growth and development. At day 20 post-infection, the size on average of the *T. pallidum*-infected organoids was significantly smaller than that of the control (*P<*0.01) (N=15 organoids from three separate bioreactors per group), and the edges of the organoids were uneven, similar to signs of disintegration; while the control group grew well (Figure 1B). To address whether *T. pallidum*-infected organoid modulates the expression of gene markers for three germ layers in neurodevelopment, i.e., endoderm, mesoderm and ectoderm, quantitative real-time PCR (qRT-PCR) was performed (N=15 organoids from three separate bioreactors per group). The qRT-PCR analysis revealed that the expressions of endodermal marker Ve-cad, KDR, and SOX17 were significantly upregulated in the *T. pallidum* group (*P* < 0.05). In contrast, expressions of the ectodermal genes MAP2, Nestin, and SOX2 were significantly decreased (*P* < 0.05) (Figure 1C). This finding was confirmed by immunofluorescent staining against TUBB3 and SOX2, which are expressed in the central nervous system from the earliest developmental stages throughout vertebrate evolution(Mercurio et al., 2019). *T. pallidum* infection resulted in brain organisational changes in neural rosette-like structures resembling the proliferative regions of the human ventricular zone where NPCs reside (Krenn et al., 2021), and caused fewer and incomplete rosette-like structures (*P* < 0.01) (Figure 1D) (N=15 organoids from three separate bioreactors per group). Cleaved caspase 3(clCASP3) staining showed that the number of apoptotic cells increased significantly following *T. pallidum* infection, but the proportion of apoptotic cells in both groups of brain organoids was very low (Figure supplement 2) (N=12 organoids, each group from three independent bioreactors). These results suggest that *T. pallidum* infection influenced the formation of neural rosette structures and affected the neurodevelopment of the brain organoid.

**Figure 1.**
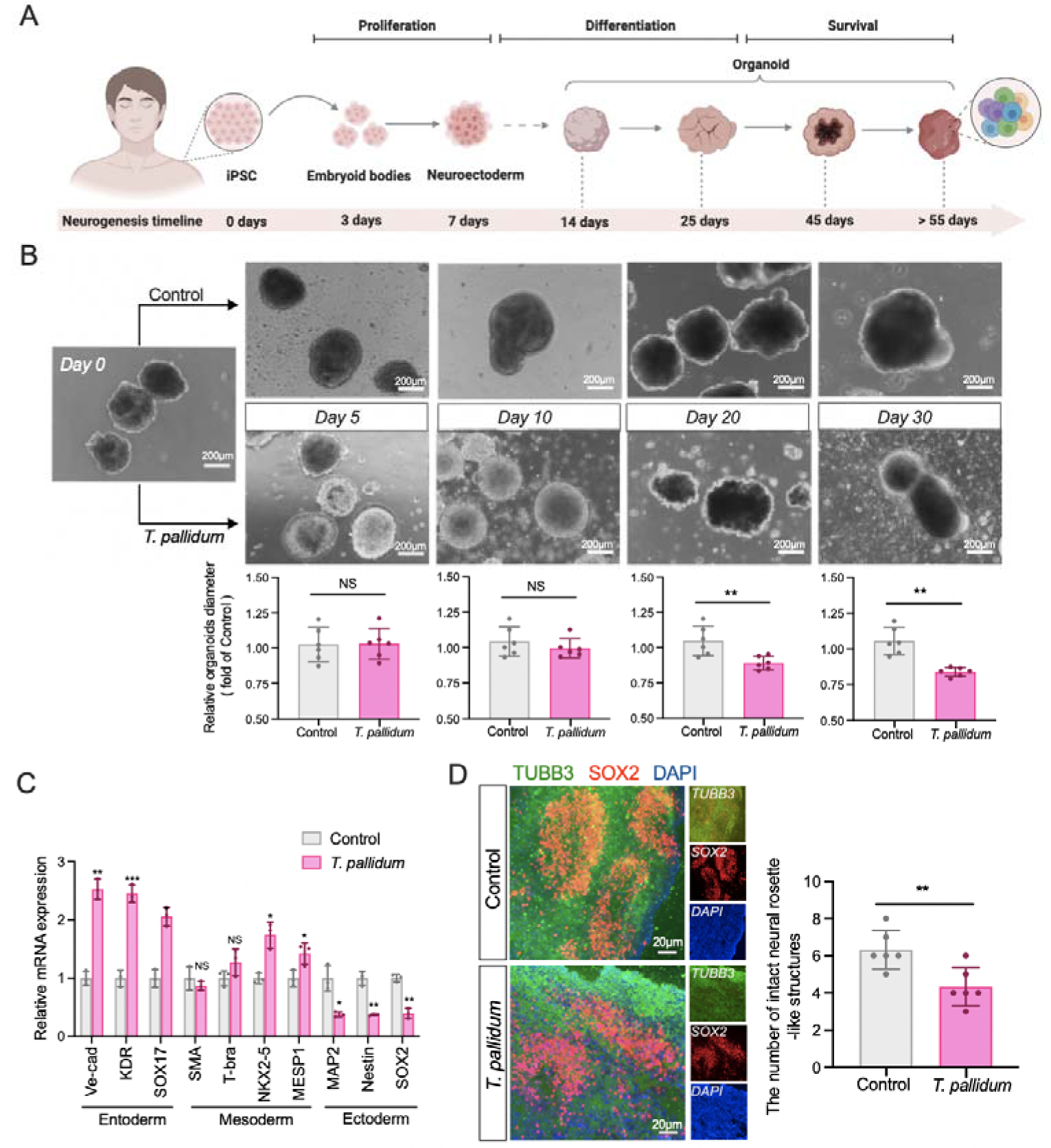
*T. pallidum* infection inhibited brain organoid neurodevelopment. (**A**) Schematic view of the methods for generating brain organoid from iPSCs. (**B**) Representative bright field images of individual brain organoids treated with *T. pallidum* over time. (**C**) Changes of the transcription levels of genes related to the three germ layers following *T. pallidum* infection in the brain organoids. (**D**) Immunofluorescent assay of the effect of *T. pallidum* infection on the brain organoid. A nonparametric *t*-test was used to evaluate the statistical differences between the two groups. (*: *P* < 0.05, **: *P* < 0.01, ***: *P* < 0.001, NS: not significant).

### scRNA-seq revealed that *T. pallidum* infection inhibited the differentiation of neural progenitor cell subcluster 1B in the brain organoid

Individual genetic markers did not address the cellular diversity of the human brain, where closely related cell classes can be identified by combining genetic signatures, while single-cell RNA sequencing allows for systematic analysis of many genes. To better understand the effects of *T. pallidum* infection on organoid development, especially cell type specification changes, scRNA-seq was performed to molecularly profile 21,504 single cells isolated from 12 organoids grown in three separate bioreactors on day 55 (after *T. pallidum* treatment for 30 days) (Figure 2A). After quality control, a mass of data from single cells of brain organoids from the *T. pallidum* (10,138 cells) and control groups (11,366 cells) were obtained. All single-cell transcriptomes underwent unsupervised clustering using *t*-distributed stochastic neighbour embedding (tSNE), yielding a total of 19 clusters (Figure 2B), which were then assigned to 13 cell types based on the known marker-gene expression enrichments (Figure 2C, Supplement file-figure supplement 3). Thereafter, we compared the regional specification in organoids between two groups, and analysed them at the single-cell level. The proportions of neural progenitor cell (NPC)1, NPC2 and neuron population were 12.5%, 1.72%, and 9.1%, respectively, following *T. pallidum* infection (*P* < 0.05), while the proportion of the neural ectoderm cell, neural epithelial cell, and undefined cells in the *T. pallidum* group were higher than that in the control group (*P* < 0.05) (Figure 2D). To identify the NPC subclusters that could be particularly vulnerable to *T. pallidum* infection, we analysed the expression of candidate genes across the NPC cell populations on brain organoids following *T. pallidum* infection. Heatmaps revealed that the key transcription factors of neural differentiation, such as HOTAIRM1, HOXB2, and HOXA2 were downregulated following *T. pallidum-*infection in the NPC1 cluster and genes regarded as neural repairing markers, such as EN2, IGFBP1 and SPRY1 were upregulated following *T. pallidum* infection (Figure 2E). However, no change in these genes was found in the NPC2 cluster between the two groups (Supplement file-figure supplement 4), illustrating that *T. pallidum* directly inhibited the development of the NPC1 cluster.

**Figure 2.**
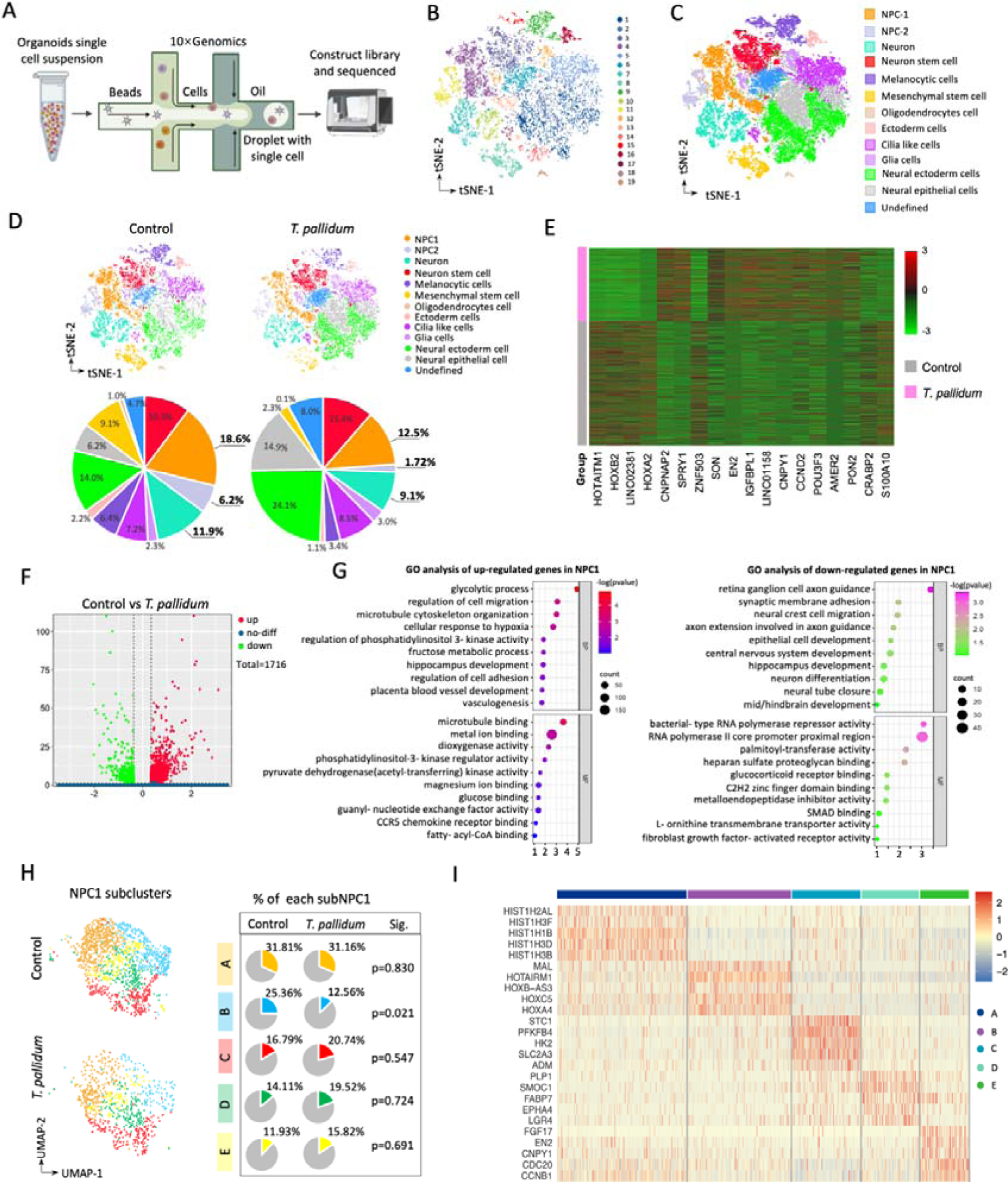
scRNA-seq revealed that *T. pallidum* infection inhibited the differentiation of neural progenitor cell subcluster 1B in the brain organoid. (**A**) Schematic overview of the 10×Genomics scRNA-seq procedure and analysis. (**B**) tSNE plot of single cells identified a total of 19 clusters. (**C**) tSNE plot of single cell depicting the separation into the 13 cell populations. (**D**) Distinguishing the tSNE plot of cell types and proportion analysis of 13 cell populations within the two groups. (**E**) Expression profile of key genes related to NPC1 and NPC2 cell populations between the two groups. Powder blue: control group; Pink: *T. pallidum* group; Pale green: NPC1; Yellow: NPC2; Red indicates high gene expression; Green indicates low gene expression. (**F**) Volcano plot of differential genes (*T. pallidum* vs control) in the NPC1 cluster. (**G**) GO analysis of the upregulated genes (left) and downregulated genes (right) in the NPC1 cluster following *T. pallidum* infection. (**H**) UMAP plot of distinguished NPC1 subclusters (left) and proportion analysis of each subcluster within two groups (right). Chi-square test was used to compare the changes in the proportion of the five small subgroups among the different groups. (**I**) Specificity marker expression of NPC1 subcluster.

To probe the effect of *T. pallidum* in the NPC1 cluster development of brain organoids, we first performed a whole-transcriptome analysis of the NPC1 cluster to discriminate any changes in the transcriptional level between the *T. pallidum* (1,290 cells) and control groups (2,114 cells) (Figure 2F). Then, Gene ontology (GO) term analysis of the upregulated and downregulated genes were identified, respectively. The upregulated genes revealed that the biological processes were mainly those related to metabolic process and blood vessel genesis, and the molecular function was closely associated with pathogenic infection. Moreover, the biological processes mainly enriched those processes associated with neural development and neurofunctional zoning, and the molecular function enriched genes promoting neural differentiation with downregulated expression (Figure 2G).

To dissect the specific cell type(s) affected by *T. pallidum* in the NPC1 population, we re-clustered and subdivided the NPC1 population into five subclusters (subNPC1 A–E) based on the high expression of general NPC1 subcluster markers (Figure 2H). The proportions of subclusters between the two groups were analysed by uniform manifold approximation and projection (UMAP) and clustering of the NPC1 population. Comparative analysis of the NPC1 subclusters between the two groups demonstrated that *T. pallidum* downregulated the number of subNPC1B population (*P* < 0.05); however, no significant difference was observed in the other subclusters. The highly expressed genes in the subNPC1B population were related to the HOX family, which is considered positively regulated for regional division and functional formation during brain development (Figure 2I). Overall, these results indicated that *T. pallidum* reduced the cell number of subNPC1B subclusters and affected brain organoid development.

### scRNA-seq revealed that *T. pallidum* infection inhibited neuronal differentiation in the brain organoid

To infer inhibition in the neuron cluster by *T. pallidum*, differential genes were grouped for volcano plot analysis (2292 cells) (Figure 3A). In the neuron population, compared with those in the control group, the upregulated and downregulated genes in the *T. pallidum* group were evaluated by GO analysis; the main functions of these upregulated genes were enriched in the items related to neuronal apoptosis and endocytosis, while the downregulated genes were enriched for cerebrum functional partition and energy metabolism in neuronal related functions, including neuron fate commitment, brain segmentation, neural tube development (Figure 3B). These results indicated that the altered genes following *T. pallidum* infection were closely linked to the neuronal dysfunctional and cerebral partition. To examine the abundance of target gene transcripts based on scRNA-seq data, eight neural-specific highly expressed genes were selected and validated. The results demonstrated that the early neural-specific transcription factors HOXA5, HOXC4, HOXC5, and NEUROG1; the positive regulatory factors during neural maturation PCP4, HOTRAIRM1 and GNG5; and the key RNA binding protein for nerve system segmentation HNRNPH1 were significantly downregulated following *T. pallidum* infection (*P <*0.01–0.001) (Figure 3C). We subsequently validated the genes by performing qRT-PCR, and found that the HOXA5, HOXC5, and HOXA4 genes of *T. pallidum* group were significantly lower than that of the control group (*P <*0.05–0.01) (Figure 3D) (N=15 organoids from three separate bioreactors per group). Immunostaining of day 55 brain organoids of neuron marker-MAP2 revealed that the neural rosette-like structure was damaged following *T. pallidum* infection (Figure 3E) (N=15 organoids from three separate bioreactors per group).

**Figure 3.**
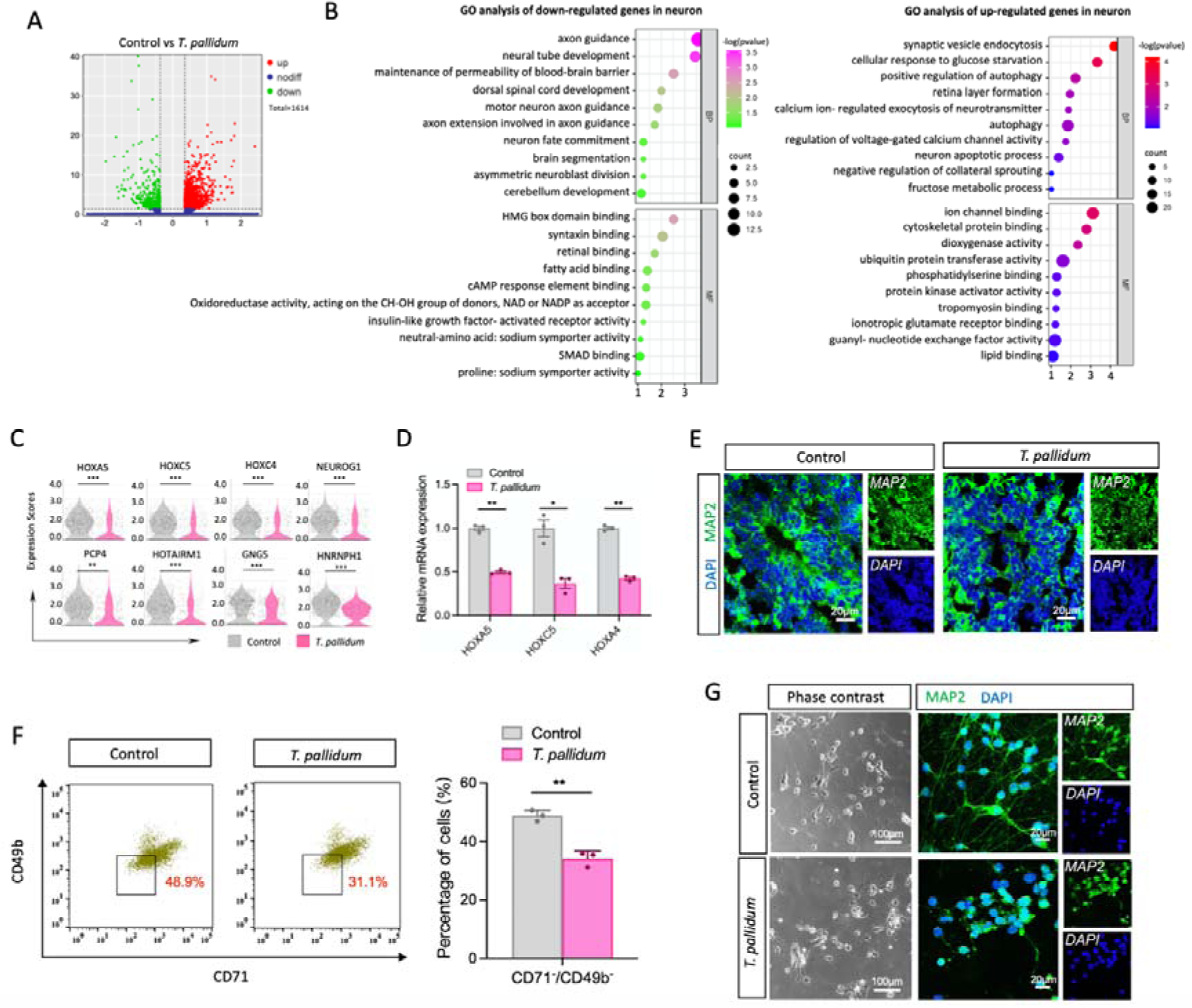
*T. pallidum* infection inhibited the differentiation of neurones in the brain organoid. (**A**) Volcano Plots of differentially expressed genes related to neurons between the two groups. (**B**) Representative GO terms of downregulated and upregulated enrichment genes related to the neurons between two groups. (**C**)Violin diagram of genes related to early neuronal development. (**D**) The effect of *T. pallidum* infection on the transcription level of genes related to the nervous system in the brain organoids. (**E**) The frozen section of brain organoids by MAP2 fluorescence staining. (**F**) Flow sorting of brain organoids and statistical analysis of the two groups using a nonparametric *t*-test. (**G**) Cell morphology and validation of CD71^−^/CD49b^−^ neuron populations. A nonparametric *t*-test was used to evaluate the statistical differences between the two groups. (*: *P* < 0.05, **: *P* < 0.01, ***, *P <*0.001, NS: not significant).

Following the confirmation that CD71 and CD49b are expressed in pluripotent stem cells, neural crest, neural stem cells but not in neurons (Menon et al., 2019), the identified difference was expended to develop novel cell selection strategies. We isolated CD71^−^/CD49b^−^ subpopulations from day 55 brain organoids (N=21 organoids from four separate bioreactors per group) differentiation using fluorescence-activated cell sorting (FACS), which revealed the proportion of CD71^−^/CD49b^−^ neurons in the *T. pallidum* group was significantly lower than that in the control group (*P* < 0.01), indicating that *T. pallidum* inhibited the normal differentiation of brain organoids into mature neurons (Figure 3F). The phase contrast images of 2-day cultured post-CD71^−^/CD49b^−^ subpopulation sorting demonstrated that the neurite of CD71^−^/CD49b^−^ neuron populations were sparse and mostly fragmented in the *T. pallidum* group, and the immunocytochemistry of neuronal markers-MAP2 increased perinuclear immunofluorescence were affected by *T. pallidum* (Figure 3G). These results indicate that *T. pallidum* inhibited the differentiation of brain organoid neurons.

### scRNA-seq revealed that *T. pallidum* infection inhibited the differentiation of the hindbrain neurons of the brain organoid

Neuron segmentation is a key step for neuronal digital reconstruction, which is essential for exploring brain circuits and understanding brain function (Li and Shen, 2022; Sugino et al., 2019). Distinct neuronal cell types acquire and maintain their identity by expressing different genes (Nelson et al., 2006). To clarify the region that was specifically inhibited by *T. pallidum*, UMAP analysis revealed the production of both unique and overlapping cell types between the two groups (Figure 4A). Based on the cerebral cortex-specific markers in each segmentation, the neuron cluster was grouped into four subclusters: hindbrain neuron, choroid plexus, noncerebral neuron, and cortical neuron (Figure 4B). Thereafter, the cell number proportions of four subclusters within the two groups were calculated; the proportion of hindbrain neuron subclusters decreased to 14.23% following *T. pallidum* infection, while the proportions of the other three subclusters showed no obvious changes between the two groups (Figure 4C). The neural-specific highly expressed genes of four subclusters were selected and validated, and the results depicted that the hindbrain neuronal markers MAGEH1, MEIS3 and USP47 were significantly decreased following *T. pallidum* infection (*P<*0.05–0.01), whereas no significant alterations were detected in the other three subclusters between the two groups (*P* > 0.05) (Figure 4D). Furthermore, immunostaining for hindbrain neuron specific markers-SHOX2 and HOXA5 in the cryosections from day 55 brain organoid revealed that the fluorescence intensity of SHOX2 and HOXA5 decreased following *T. pallidum* infection (Figure 4E) (N=15 organoids from three separate bioreactors per group). Overall, these results suggest that *T. pallidum* inhibited the differentiation of hindbrain neurons in the brain organoids.

**Figure 4.**
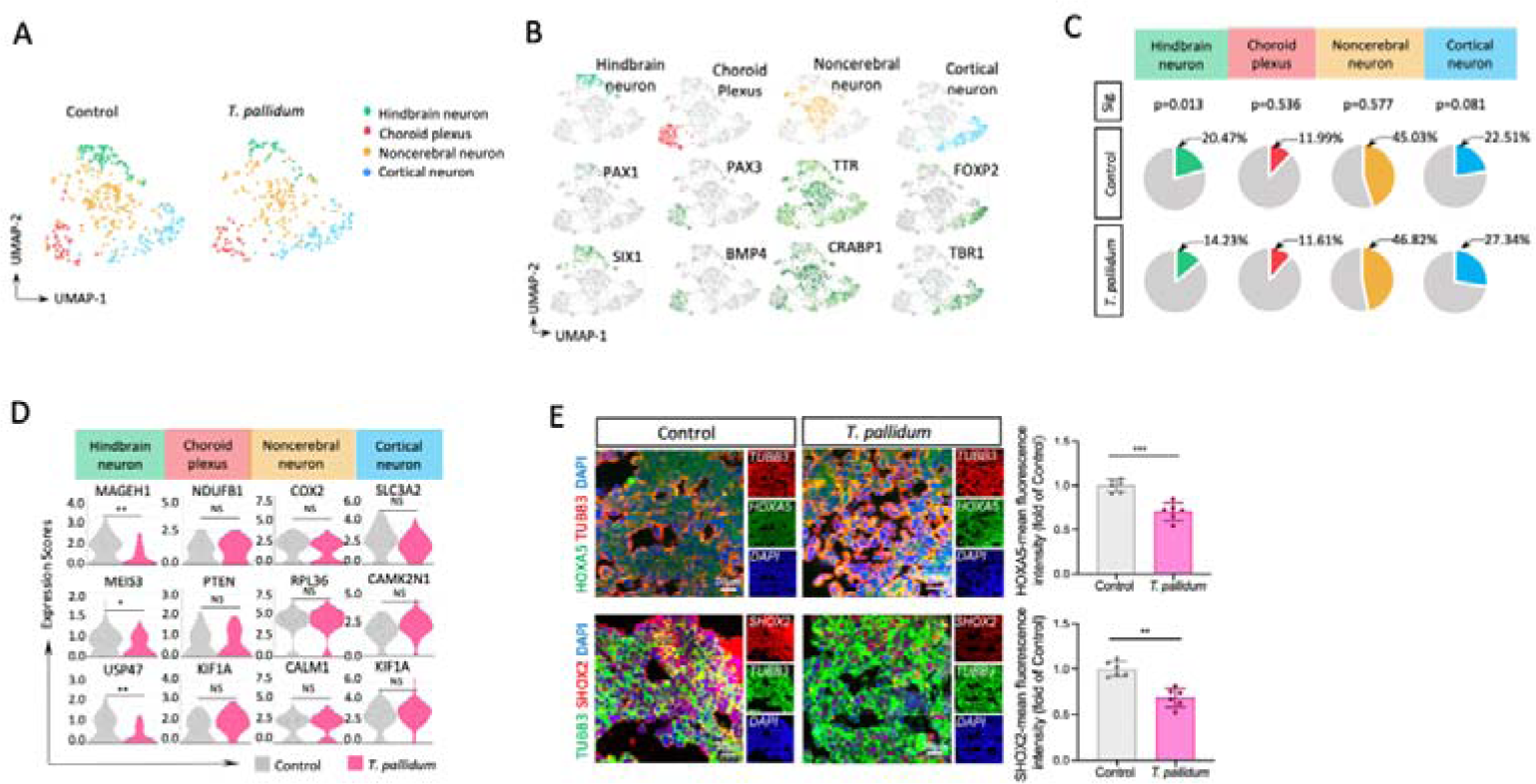
scRNA-seq revealed that *T. pallidum* infection inhibited the differentiation of the hindbrain neurons in the brain organoid. (**A**) UMAP embedding plots comparing the distribution of the brain organoids within the two groups. (**B**) UMAP embedding of cell subset in each region of the neurons calibrated by marker genes. (**C**) Subcluster cell number as a percentage of the neuron cluster cell number. The Chi-square test was used to compare the differences in the percentages among different cell subsets. (**D**) Violin diagram of the relevant marker genes in each neuronal region. (**E**) Immunohistochemistry of the brain organoids validating the expression of hindbrain neuron markers, including HOXA5 and SHOX2. A nonparametric *t*-test was used to compare the differences between the two groups (*: *P* < 0.05, **: *P* < 0.01, NS: not significant).

### scRNA-seq revealed the mechanism of *T. pallidum* affecting neurodevelopment in the brain organoids

To study the differentiation of subNPC1B into hindbrain neuron cluster and the corresponding gene expression, we selected HES4 and NEUROGD1 to construct a new trajectory corresponding to two distinct cell fates. The root of the trajectory was mainly populated by HES4 for subNPC1B subcluster, while the termini of the tree was populated by NEUROGD1 for the hindbrain neuron subcluster (Figure 5A). Thereafter, we assessed the expression of the genes-regulated trajectory of the subNPC1B-hindbrain neuron. ASPM, CCNB1, TPX2, UBE2C, TOP2A, MKI67, CENPF, PTTG1 and HMGN2 expression, which were the top genes in subNPC1B, were markedly reduced from the root to the termini. SNCG, S100A10, NEUROD1, RGS10, STMN2, PPP1R17, BASP1, NEFL, CALM1, DCX, TMSB10, TUBB3, TMSB4X, STMN4, C11orf96, TUBA1A, RTN1, SYT4, and SST expression, which were the top genes in the hindbrain neuron subcluster, were lowly expressed at the early stage of the pseudotime axis, and gradually increased until the end of the pseudotime axis (Figure 5B). These results further confirmed that the differentiation direction is from subNPC1B to the hindbrain neuron.

**Figure 5.**
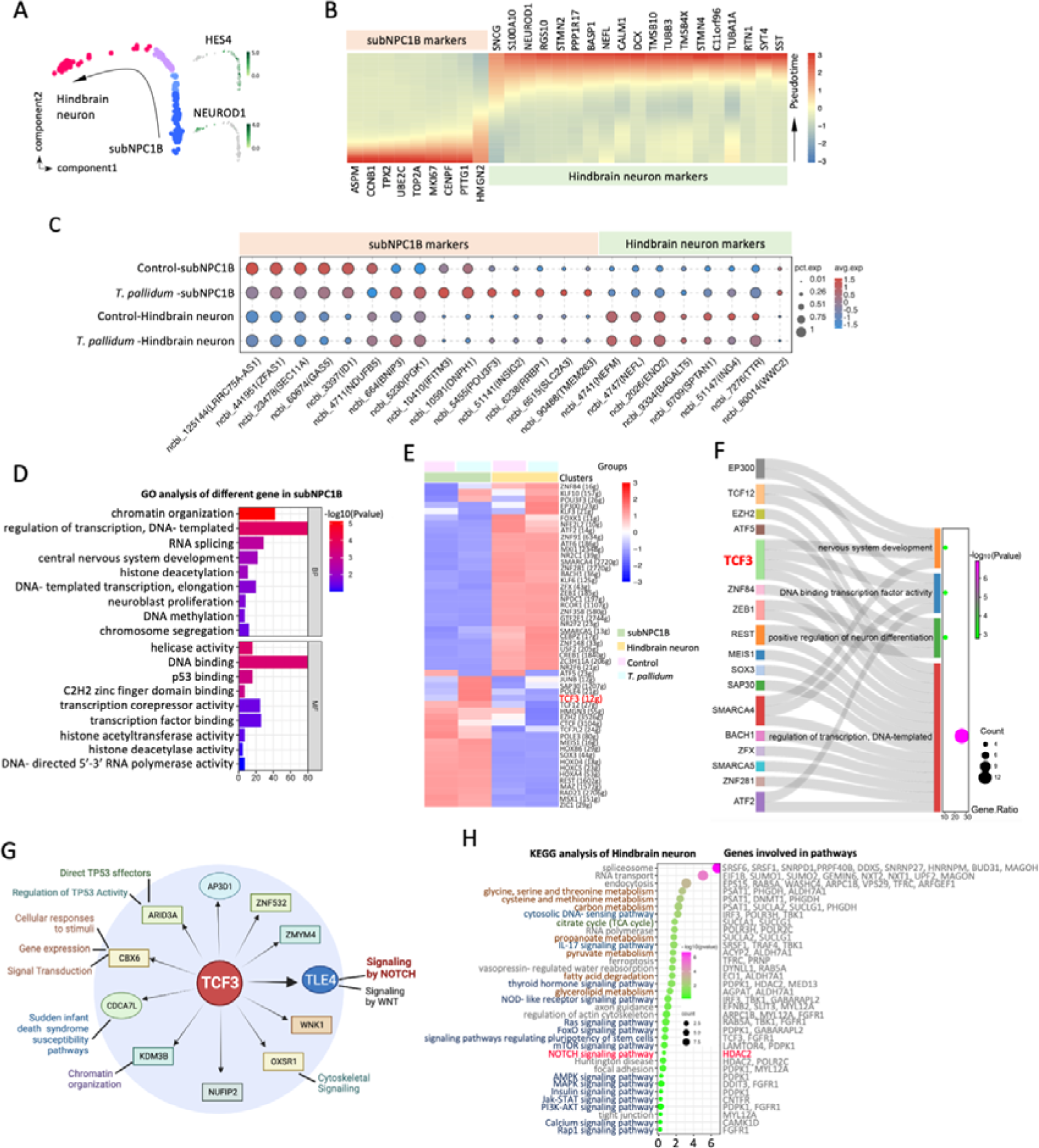
scRNA-seq revealed the mechanism of *T. pallidum* affecting neurodevelopment in the brain organoids. (**A**) Left panel: temporal analysis of subNPC1B-hindbrain neuron by Monocle 2, with subNPC1B subgroup in blue and hindbrain neuron subgroup in magenta. Right: HES4 localises to subNPC1B and NEUROGD1 localises to the hindbrain neuron. (**B**) Heatmap of differentially expressed transcripts along pseudotime. Cells are ordered by cell type and pseudotime. (**C**) Bubble difference diagram of subNPC1B and hindbrain neuron marker genes before and after *T. pallidum* infection. (**D**) GO analysis of the total number of differential genes in the subNPC1B subgroup. (**E**) The differential expression of transcription factors between the *T. pallidum* and control group by SCENIC analysis. (**F**) Sankey diagram showing the correspondence between differential transcription factors and neural differentiation and development. (**G**)Extended analysis of TCF3. (**H**) KEGG enrichment pathway analysis of hindbrain neuron subgroup.

To determine whether *T. pallidum* inhibited the differentiation of hindbrain neurons by disturbing subNPC1B-hindbrain neuron differentiation axis or by directly affecting the hindbrain neuron, subNPC1B and hindbrain neuron markers were displayed using dot shapes (Figure 5C). The subNPC1B specific markers-LRRC75A-AS1, ZFAS1, SEC11A, GAS5, ID1, and NDUFB5 were decreased following *T. pallidum* infection; the inhibitors markers-BNIP3, PGK1 and IFITM3 during the NPC differentiation were increased; the early NPC markers-POU3F3 and INSIG2 also increased in the *T. pallidum* group; however, there was no difference in the expression of the hindbrain neuronal markers-NEFM, NEFL, ENO2, B4ALT5, SPTAN1, and TTR in either the *T. pallidum* group or control group, indicating that the decreased number of hindbrain neurons was attributed to the disturbance of the subNPC1B-hindbrain neuron differentiation axis.

To uncover the mechanism underlying the subNPC1B-hindbrain neuron differentiation process, GO analysis of the total differential genes in subNPC1B demonstrated that the GO terms were mainly enriched in transcription regulation (Figure 5D), suggesting that the transcriptional regulatory network mediated by the transcription factors of subNPC1B could play a major role during the inhibition of the subNPC1B-hindbrain neuron differentiation process by *T. pallidum*. Next, the single-cell regulatory network inference and clustering (SCENIC) analysis for subNPC1B subcluster, which was performed to assess the differences in the transcriptional activity of the transcription factors between the two groups, revealed that the transcriptional activity of 52 genes significant changed after *T. pallidum* infection (Figure 5E). Furthermore, GO analyses demonstrated that 27 transcription factors were significantly enriched in the key pathways of neural differentiation and development in response to nervous system development, positive regulation of sequence-specific DNA-binding transcription factor activity, positive regulation of neuronal differentiation, and DNA templated transcription regulation. Remarkably, transcription factor 3 (TCF3) is the sole transcription factor present in all four terms simultaneously (Figure 5F), speculating that TCF3 is the key transcription factor for the inhibition of subNPC1B-hindbrain neuron differentiation caused by *T. pallidum*. Furthermore, we explored TCF3 function which could be extended to known TCF3-specific downstream factors and pathway activities (Figure 5G), Kyoto Encyclopedia of Genes and Genomes (KEGG) pathway enrichment analysis subsequently performed on the hindbrain neuron subcluster (Figure 5H), and found that they were in the common notch signaling pathway. Therefore, we speculate that TCF3 and notch signalling pathway could play important roles in the inhibition of subNPC1B-hindbrain neuronal differentiation axis by *T. pallidum*, warranting further study to confirm these findings.

## Discussion

Infants born to mothers with *T. pallidum-*infection and who had a positive syphilis serologic test during pregnancy have a high risk of developing neurodevelopmental disorders (Auriti et al., 2022; Verghese et al., 2018), with varied clinical manifestations, such as bulging fontanelle, seizures, and cranial nerve palsies (Mattei et al., 2012; Woods, 2009). Owing to ethical and practical challenges, guinea pig and rabbit animal models have been widely employed to study the pathogenesis of congenital syphilis (Wicher and Wicher, 2001); however, neither neuroinvasive nor congenital syphilis has been well explained (Tantalo et al., 2005). In the study, we evaluated iPSC-derived brain organoids for modelling to reveal the pathogenesis of neurodevelopmental disorders associated with congenital syphilis. The results showed that *T. pallidum* infection inhibited brain organoid neurodevelopment. This conclusion is supported by the results showing brain organoids infected with *T. pallidum* demonstrated significant decrease in the overall organoid size and interfered with the formation of neural rosette-like structure of brain organoids. Combined with scRNA-seq analysis, it clarified that *T. pallidum* inhibited the differentiation of hindbrain neurons through the suppression of subNPC1B in brain organoids. Furthermore, we found that TCF3 may be a key transcription factor associated with the inhibition of subNPC1B-hindbrain neuron axis by *T. pallidum*.

As brain organoids reflect developmental stages during pregnancy and the early postnatal, they are currently the most relevant tools for research on neurodevelopmental disorders (Andrews and Kriegstein, 2022), helping to discover novel biological information on the early stages of neurodevelopment and disease progression, as well as the ability to manipulate these processes *in vitro* (Trujillo and Muotri, 2018). Tonni et al. reported that a routine third-trimester scan of pregnant woman with primary syphilis, which revealed foetal growth restriction associated with absent foetal movement, a pathologic neuroscan characterised by cortical calcifications, and a transabdominal neuroscan was able to detect hyperechogenicity involving both choroid plexuses of the lateral ventricles as well as the falx cerebri, with smooth without recognisable gyri and sulci (Tonni et al., 2022). The imaging features and the neurodevelopmental outcome in the fetal brain were verified by organoids infected with *T. pallidum* inducing a significant decrease in the overall organoid size and interfering with the formation of neural rosette-like structure of brain organoids in our study. Previous studies have shown that Zika virus-induced cellular death and reduced size of cortical organoids are associated with the clinical microcephaly phenotype (Nowakowski et al., 2016; Qian et al., 2016; Watanabe et al., 2017). Whether *T. pallidum* induced the decrease in overall organoid size and disruption of neural rosette-like structure in the brain organoids is related to the clinical microcephaly phenotype required further confirmation in the infection animals.

Uterine environmental pollution taking place in early brain development could have long-lasting consequences for adult brain functioning (Meyer et al., 2007). Several studies demonstrated that the development of the central nervous system was blocked at an immature stage following Zika virus infection (Hughes et al., 2016; Li et al., 2016; Zhang et al., 2019), preferentially targeting and hindering the growth of neural precursor cells in the monolayer and cortical brain organoid culture systems (Zhang et al., 2019). Notably, similar results were observed in our study: the proportion of neuroectodermal cells, neuroepithelial cells, neural stem cells, and other cells in the early stages of neurodevelopment of *T. pallidum*-infected organoids were significantly higher than that the control group, suggesting that organoids infected with *T. pallidum* retained neural precursor cells in the developmental stage and inhibited their differentiation in brain organoids towards maturity. Besides, NPCs appear to be the most susceptible cells of the central nervous system to Zika virus infection (Rubio-Hernández et al., 2023). Zika virus can directly infect the NPC and disrupt neural development, resulting in reduced organoid growth and overall size, thereby recapitulating microcephaly observed in foetuses and infants exposed to the virus (Watanabe et al., 2017). In this study, the proportion of NPC and neuron population decreased following *T. pallidum* infection, suggesting that *T. pallidum* inhibited the differentiation of immature cells into NPCs in the brain organoids, resulting in the number of NPC (especially subNPC1B), and subsequently led to a decrease in the number of neurons. Moreover, 40% of children with congenital syphilis have similar mental disorders to those in congenital Zika virus infection (Carrier and Haughton, 2019).

The vertebrate hindbrain contains a complex network of dedicated neural circuits that play an essential role in controlling many physiological processes and behaviours, including those related to the cerebellum, pons, and medulla oblongata(Shoja et al., 2018). Patients with pontocerebellar hypoplasia represent the less severe end of the spectrum with early hyperreflexia, developmental delay, and feeding problems, eventually developing spasticity and involuntary movements in childhood, while some patients represent the severe end of the spectrum characterised by polyhydramnios, severe hyperreflexia, contracture, and early death from central respiratory failure. Patients with pontocerebellar hypoplasia develop microcephaly at birth or over time after birth (Doherty et al., 2013). The hindbrain also contains a wide variety of inter- and relay neurons, including a network of reticulospinal neurons that regulate alertness, sleep, posture, and fine-grained locomotor activities(Krumlauf and Wilkinson, 2021; Valakh et al., 2023). Our scRNA-seq revealed that the number of hindbrain neurons significantly decreased following *T. pallidum* infection, which supports the fact that *T. pallidum* infection causes systemic paresis, progressive dementia, microcephaly and other nervous system related symptoms. HOX family genes are key factors regulating early hindbrain development and segmentation (Frohman et al., 1990; McNulty et al., 2005). SHOX2 is required for vestibular statoacoustic neuronal development with nerve fibres mostly terminating in the pons and medulla oblongata (hindbrain regions) (Laureano et al., 2022). Immunofluorescence staining of HOXA5 and SHOX2 markers demonstrated that the fluorescence intensity of SHOX2 and HOXA5 decreased following *T. pallidum* infection, thereby revealing that *T. pallidum* inhibited the development and maturation of hindbrain neurons on organoids, which was again confirmed by the scRNA-seq analysis results. Moreover, our scRNA-seq analysis revealed that *T. pallidum* inhibited the hindbrain neuron cell number through the suppression of subNPC1B population in organoids, and found that TCF3 and notch signalling pathway may play important roles in the inhibition of subNPC1B-hindbrain neuron differentiation axis by *T. pallidum*. However, in-depth exploration of the mechanism of *T. pallidum*-inhibited hindbrain neurogenesis is warranted for further understanding the pathogenesis and early intervention of congenital neurodevelopmental impairment.

Nevertheless, certain limitations remain to be addressed and could likely be the topic of future studies. First, although several recent protocols have made use of growth factors to promote further neuronal maturation and survival (Lucke-Wold et al., 2018), the organoid culture scheme needs to be further improved owing to the lower percentage of mature neurons and the challenge of cell necrosis within the organoids at this stage in day 55 organoids. Second, only changes in the appearance, morphology, and cell type of the organoids infected with *T. pallidum* were observed in this study, while research on physiological function was still blank. Third, further experiments are warranted to confirm that *T. pallidum* restricts the neurodevelopment of organoids by inhibiting the subNPC1B-hindbrain neuron differentiation axis, which is limited to bioinformatics analysis. Finally, future animal experiments could allow a more comprehensive understanding of the mechanisms discovered in organoid studies.

In summary, even though the current organoid systems demonstrated limitations and require additional optimisation for better use in disease modelling, they have opened up important novel avenues for related research on congenital neurodevelopmental impairment. Based on the bioinformatics analysis of scRNA-seq, we found that *T. pallidum* could inhibit the differentiation of subNPC1B in brain organoids, thereby reducing the differentiation from subNPC1B to hindbrain neurons, and ultimately affecting the development and maturation of hindbrain neurons during pregnancy. The TCF3 and notch signalling pathway could play important roles in this process, which needs further experimental verification. This is the first report to study the effects of *T. pallidum* on the neural development of an iPSC-derived brain organoid model, which could serve as a critical benchmark for studying the mechanism of congenital neurodevelopmental impairment in the future.

## Materials and methods

### Preparation of *T. pallidum*

The *T. pallidum* Nichols strain was kindly provided by Lorenzo Giacani, PhD (University of Washington, Seattle, WA, USA) and propagated in male New Zealand rabbits as described previously(Tong et al., 2017). *T. pallidum* was mixed with the organoids maturation stage medium and thereafter used to treat the organoids.

### iPSC culture and brain organoids generation

iPSC was purchased from the American Type Culture Collection, which were derived from fibroblasts that were reprogrammed by the ectopic expression of four specific genes (OCT4, SOX2, KLF4, and MYC)(Yang et al., 2022). Brain organoids were generated from iPSC using established protocols as previously described(Lin et al., 2018; Madhavan et al., 2018), but slightly modified. Briefly, iPSC colonies were cultured in a 6-well plate coated with Matrigel (Corning, 354277) in the mTeSR™ medium (STEMCELL Technologies, 85851) until the cells reached 90–100% confluence and dissociated into single cells with Accutase (Gibco, A1110501). Thereafter, the iPSCs were transferred to a 6-well plate with Ultra-Low Attachment surface polystyrene (Corning, REF3471) in StemFlex™ DMEM/F12 Medium (Gibco, A33493) containing with 20% KnockOut Serum (Gibco, 10828028), 1×NEAA (Gibco, 11140-050), 1×GlutaMAX-1 (Gibco, 35050-061), 1×β-Mercaptoethanol (Invitrogen, 21985-023), 5 mM Dorsomorphin (Sigma, P5499), 10 mM SB-431542 (Sigma, S4317), and 1% Penicillin-Streptomycin (Gibco, 15140-0122); 10 μM Rock inhibitor (MCE, Y-27632) was added to the medium after the first 3 days and then the medium was half changed every other day. At day 7, the old medium was removed and replaced with a differentiation medium consisting of Neurobasal-A-Medium (Gibco, 10888-022) containing 1×B27 (Gibco, 12587010), 1×GlutaMAX-1, 20ng/ mL FGF2 (R&D Systems, 233-FB), 10 ng/mL EGF (R&D Systems, 236-EG), 1×β-Mercaptoethanol, and 1% Penicillin-Streptomycin. On day 25, the differentiation medium was exchanged with a maturation medium consisting of Neurobasal-A-Medium, 20ng/mL BDNF (R&D Systems, 248-BDB), 20 ng/mL GDNF (R&D Systems, 212-GD), 20 ng/mL NT-3 (R&D Systems, 267-N3), 10 ng/mL EGF, and 1% Penicillin-Streptomycin. The medium was half changed every other day until 55 days.

### Immunohistochemistry

For immunohistochemistry of the brain organoids, organoids were processed at day 55 as previously described(Qian et al., 2016). Briefly, whole organoids were fixed in 4% paraformaldehyde for 45–60 min with slow shaking at 25 C. The organoids were washed three times with phosphate buffer solution (PBS) and then incubated in 30% sucrose solution overnight at 4 C. The organoids were embedded in plastic cryomolds (Tissue Tek) and filled with O.C.T cryoembedding medium (Sakura, 4583). The embedded tissue was frozen on dry ice and then stored at –80 C until cryostat sectioning. Frozen organoid tissue was sliced into 15-µm sections using a microtome cryostat (MICROM International GmbH, HM550) at −15 to −20CC, and collected using superfrost Ultra Plus slides. For immunostaining, the organoid sections were quickly washed with PBS to eliminate any residual O.C.T, and then blocked with blocking permeabilising solution (0.3% Triton-X, 0.2% Tween-20, 0.1% bull serum albumin and 10% normal goat serum in PBS) for 1 h at room temperature. Thereafter, sections were incubated overnight at 4CC in an antibody diluent (1.1% Triton-X and 2% normal goat serum in PBS) containing primary antibodies anti-TUBB3 (mouse, 1:1000, Sigma-Aldrich, T8660; rabbit, 1:300, Abcam, ab18207), anti-SOX2 (rabbit, 1:300, Abcam, ab92494), anti-MAP2 (rabbit, 1:200, Abcam, ab32454), anti-cleaved caspase 3 (rabbit, 1:100, Cell Signaling Technology, 9661S), anti-SHOX2 (rabbit, 1:100, Affinit, DF14735), and anti-HOXA5 (mouse, 1:100, Santa cruz, sc-365784). On the following day, sections were rinsed three times in PBST (PBS containing 0.05% Tween 20) before incubation for 1 h at room temperature in secondary antibody solution, which contained Alexa Fluor 488-conjugated anti-mouse antibody (goat, 1:1000, Thermo Fisher, A11001) and Alexa Fluor 647-conjugated anti-rabbit antibody (goat, 1:1000, Thermo Fisher, A-21244). Finally, sections were washed with PBST and stained with VECTASHIELD^®^ Antifade Mounting Medium with DAPI (Vector laboratories, H1200). Stained organoid cryosections were imaged using a confocal laser scanning microscope (Zeiss, LSM 700).

For immunohistochemistry of neurons, following culture on Millicell glass coverslips, neurons obtained by flow sorting were washed with PBS three times and fixed with 4% paraformaldehyde for 15 min. After fixation, the neurons were washed with PBS and permeabilised with blocking solution containing 10% normal goat serum, 2% bull serum albumin, and 0.05% Triton X-100 in PBS for 1 h at room temperature. Then neurons were incubated with anti-MAP2 (rabbit, 1:200, Abcam, ab32454) overnight at 4 C and Alexa Fluor 647-conjugated anti-rabbit antibody (goat, 1:1000, Thermo Fisher, A-21244) for 1 h at room temperature, and washed three times with PBS before mounting on the glass slide.

### Flow cytometric screening and candidate validation

Neuron cells obtained from brain organoids were screened for cell surface molecule expression using an ultra-high-speed flow cytometry sorter (Beckman, MoFlo Astrios EQS) as previously reported(Menon et al., 2019). Organoids were dissociated into single cell, and then resuspended in PBS with 5% KnockOut Serum and stained using CD71-PE (1:100, Bioscience, 12-0719-42) and CD49b-APC (1:100, Bioscience, 17-0500-42). Antibodies were incubated for 30 min at 4 °C and the cells were subsequently washed twice in PBS with 5% KnockOut Serum. For flow cytometric analysis, single cell suspensions were dispensed for sorting and then culture for subsequent observation and validation. Data analyses were performed using FlowJo™ version 10.7.1 (Becton, Dickinson and Company).

### RNA isolation and qRT-PCR

For each *T. pallidum* or control group, the organoids were collected at day 55 in 2 mL RNAse-free tubes and chilled on ice throughout the procedure. Brain organoids were homogenised in Trizol (Invitrogen, 1596026) and processed according to the manufacturer’s instructions. cDNA synthesis was performed using a TransScript All-in-One First-Strand cDNA Synthesis SuperMix for qPCR kit (Invitrogen, AT341-01) according to the manufacturer protocols. qPCR quantified cDNA via SYBR^®^ Premix Ex Taq™ II (Takara, RR82LR). Each reaction was run in triplicate and analysed following the ΔΔCt value using glyceraldehyde-3-phosphate dehydrogenase (GAPDH) as a reference gene. Data were presented as expression level (2^-ΔΔCt^) relative to GAPDH. The primer pairs are listed in Supplementary Table.

### Preparation of single-cell suspensions and scRNA-seq of brain organoids

For preparation of single-cell suspensions, the organoids were harvested and dissociated into single cells using Accutase. Single cell was resuspended with cold DMEM/F-12 medium (Gibco, C11330500BT), and counted by Trypan Blue to determine the live cell concentration. Living cell rate was preferably above 90%, and then diluted for an appropriate concentration to obtain approximately 6000 cells per lane of a 10×microfluidic chip device; 30000–50000 cells were needed to generate the scRNA-seq. Single cell sequencing experiments were conducted using the 10×Genomics Chromium following the manufacturer’s instructions by GENE DENOVO, Guangzhou, China. A detailed description of the scRNA-seq protocol is available in the experiment flow of the Supplementary Protocol.

### scRNA-seq data analysis

10× Genomics Cell Ranger software (version 3.1.0) was employed to convert raw BCL files to FASTQ files, alignment, and counts quantification. These FASTQ files were aligned to the human reference genome and transcriptome (GCF_000001405.38_GRCh38.p12). The cell by gene matrices for each sample were individually imported to Seurat (v3.1.1)(Butler et al., 2018) for downstream analysis. Cluster markers were interpreted and assigned cluster identity by using known literature cell type annotations identifiable details, and the organoids were divided into 13 cell populations using the tSNE algorithm in the R language. Subclustering within the NPC and neuron populations was performed by selecting and clustering the cells from the NPC and neuronal clusters, respectively, and repeating the clustering procedure with batch effect correction. The resulting distance matrices of NPC and neuron clusters were used as input for generating UMAP embeddings. After subclustering, the NPC was identified and subclustered into NPC1 and NPC2; NPC1 were ulteriorly subclustered and generated into subNPC1A, subNPC1B, subNPC1C, subNPC1D, and subNPC1E. Neuron populations were identified and subclustered to generate four clusters: hindbrain neuron, choroid plexus, noncerebral neuron, and cortical neuron.

Differentially expressed genes were identified using the SingleR R package(Satija et al., 2015) and cut-off of |log fold-change| > 0.25 (Bonferroni adjusted *p*-value < 0.05). Thereafter, the violin plot of differentially expressed genes was displayed using the ‘ggplot2’ R package, and the heatmap of differentially expressed genes was plotted using the ‘heatmap’ R package(Saliba et al., 2016).

To identify disturbed biological functions and understand the importance of differentially expressed genes following *T. pallidum* infection on organoids, GO classification was performed, which included the following categories: biological process and molecular functions. GO functional enrichment analysis was conducted for identifying differentially expressed genes using Database for Annotation, Visualization and Integrated Discovery(Sherman et al., 2022; Zhang et al., 2016) with default parameters. KEGG pathway enrichment analysis identified significantly enriched metabolic pathways or signal transduction pathways in the hindbrain neuron clusters. The distribution of expression of each cluster was demonstrated by using a dot plot.

To estimate RNA velocity, spliced and unspliced transcripts were enumerated using the scVelo (v0.2.4)(Bergen et al., 2020). We performed pseudotemporal trajectory inference analysis on the subNPC1B and hindbrain neurons of human brain organoids utilising Monocle2 using the DDR-Tree and default parameter.

SCENIC was used to analyse the transcription factors for identifying gene regulatory networks as previously described(Aibar et al., 2017; Lin et al., 2021a). Briefly, we predicted the gene regulatory networks and transcription factors using SCENIC (pySCENIC, v.11.2), which consisted of the following three steps: establishment of co-expression modules by Random Forest, identification of direct relationship using motif analysis via RcisTarget, and calculation of the regulon activity score with Area Under the recovery Curve (Lin et al., 2021b). The Sankey diagram was created using SankeyMATIC (https://sankeymatic.com/) (Zhang et al., 2023), which was used to characterize the interactions between differential transcription factors and neural differentiation and development. The TCF3 was identified from the subNPC1B subsets in the organoids from single-cell transcripts.

### Data visualisation and statistical analysis

All the statistical analyses and data visualisation were performed under the R statistical computing environment (v3.6.3). Heatmaps of gene expression were produced using pheatmap (v1.0.12). Visualisation of the three-dimensional data embeddings were performed using the rgl package (v0.100.54). All other data visualisations were produced using functionalities provided in Seurat and the ggplot2 package (v3.3.0). The Chi-square test was used to compare the percentage differences among different groups. One-way analysis of variance with Tukey’s multiple comparisons tests were used for multiple group comparisons. A nonparametric *t*-test was used to perform the statistical analysis of the differences in the mRNA expression levels and mean fluorescence intensity in the Prism program (v5.0). Data are presented as mean ± standard error of mean. *P* < 0.05 was considered statistically significant.

## Acknowledgements

This work was funded by the National Natural Science Foundation of China (grant number 82272370 and 81973104 to T.-C. Y., and grant number 82001600 to Y. L.), Science and Technology Program of Xiamen City, Fujian Province, China (grant number 3502Z20224036 to T.-C. Y.), the Natural Science Foundation of Fujian Province, China (grant number 2021J02055 to T.-C. Y., and grant number 2020J05292 to Y. L., grant number 2021J01073 to M.-L. T.). The funders played no role in the study design, data collection, or analyses, the decision to publish, or manuscript preparation.

## Author contributions

Q.-Y. X., Conceptualisation, Data curation, Software, Formal analysis, Validation, Investigation, Visualisation, Methodology, Writing – original draft, Writing – review and editing; Y.-J. W., Data curation, Software, Methodology; Y. H., Investigation, Visualisation, Methodology; X.-Q. Z., Data curation, Software, Methodology; M.-L. T., Resources, Formal analysis, Supervision, Funding acquisition, Validation; Y. L., Conceptualisation, Data curation, Software, Formal analysis, Validation, Investigation, Funding acquisition, Visualisation; T.-C. Y., Conceptualisation, Funding acquisition, Writing – original draft, Project administration, Writing – review and editing

## Data and materials availability

The raw scRNA-sequence data reported in this paper have been deposited in the Genome Sequence Archive (Genomics, Proteomics & Bioinformatics 2021) in National Genomics Data Center (Nucleic Acids Res 2022), China National Center for Bioinformation / Beijing Institute of Genomics, Chinese Academy of Sciences (GSA-Human: HRA004501) that are publicly accessible at https://ngdc.cncb.ac.cn/gsa-human. All data needed to evaluate the conclusions in the paper are present in the paper and/or the Supplementary Materials.

## References

Aibar S, González-Blas CB, Moerman T, Huynh-Thu VA, Imrichova H, Hulselmans G, Rambow F, Marine JC, Geurts P, Aerts J, van den Oord J, Atak ZK, Wouters J, Aerts S. 2017. SCENIC: single-cell regulatory network inference and clustering. Nat Methods 14: 1083–1086.

Andrews MG, Kriegstein, AR. 2022. Challenges of Organoid Research. Annu Rev Neurosci 45, 23–39.

Auriti C, Bucci S, De Rose DU, Coltella L, Santisi A, Martini L, Maddaloni C, Bersani I, Lozzi S, Campi F, Pacifico C, Balestri M, Longo D, Grimaldi T. 2022. Maternal-Fetal Infections (Cytomegalovirus, Toxoplasma, Syphilis): Short-Term and Long-Term Neurodevelopmental Outcomes in Children Infected and Uninfected at Birth. Pathogens 11: 1278.

Bergen V, Lange M, Peidli S, Wolf FA, Theis FJ. 2020. Generalizing RNA velocity to transient cell states through dynamical modeling. Nat Biotechnol 38: 1408–1414.

Butler A, Hoffman P, Smibert P, Papalexi E, Satija R. 2018. Integrating single-cell transcriptomic data across different conditions, technologies, and species. Nat Biotechnol 36: 411–420.

Camp JG, Badsha F, Florio M, Kanton S, Gerber T, Wilsch-Bräuninger M, Lewitus E, Sykes A, Hevers W, Lancaster M, Knoblich JA, Lachmann R, Pääbo S, Huttner WB, Treutlein B. 2015. Human cerebral organoids recapitulate gene expression programs of fetal neocortex development. Proc Natl Acad Sci U S A 112: 15672–15677.

Carrier J, Haughton V. 2019. Congenital Syphilis: A Challenging Case for NICU Clinicians. Neonatal Netw 38: 170–177.

David M, Hcini N, Mandelbrot L, Sibiude J, Picone O. 2022. Fetal and neonatal abnormalities due to congenital syphilis: A literature review. Prenat Diagn 42: 643–655.

Doherty D, Millen KJ, Barkovich AJ. 2013. Midbrain and hindbrain malformations: advances in clinical diagnosis, imaging, and genetics. Lancet Neurol 12: 381–393.

Fortin O, Mulkey SB. 2023. Neurodevelopmental outcomes in congenital and perinatal infections. Curr Opin Infect Dis 36: 405–413.

Frohman MA, Boyle M, Martin GR. 1990. Isolation of the mouse Hox-2.9 gene; analysis of embryonic expression suggests that positional information along the anterior-posterior axis is specified by mesoderm. Development 110: 589–607.

Grosse SD, Dollard SC, Ortega-Sanchez IR. 2021. Economic assessments of the burden of congenital cytomegalovirus infection and the cost-effectiveness of prevention strategies. Semin Perinatol 45: 151393.

Hughes BW, Addanki KC, Sriskanda AN, McLean E, Bagasra O. 2016. Infectivity of Immature Neurons to Zika Virus: A Link to Congenital Zika Syndrome. EBioMedicine 10: 65–70.

Kanton S, Boyle MJ, He Z, Santel M, Weigert A, Sanchís-Calleja F, Guijarro P, Sidow L, Fleck JS, Han D, Qian Z, Heide M, Huttner WB, Khaitovich P, Pääbo S, Treutlein B, Camp JG. 2019. Organoid single-cell genomic atlas uncovers human-specific features of brain development. Nature 574: 418–422.

Kim J, Koo BK, Yoon KJ. 2019. Modeling Host-Virus Interactions in Viral Infectious Diseases Using Stem-Cell-Derived Systems and CRISPR/Cas9 Technology. Viruses 11.

Krenn V, Bosone C, Burkard TR, Spanier J, Kalinke U, Calistri A, Salata C, Rilo Christoff R, Pestana Garcez P, Mirazimi A, Knoblich JA. 2021. Organoid modeling of Zika and herpes simplex virus 1 infections reveals virus-specific responses leading to microcephaly. Cell Stem Cell 28: 1362–1379.e1367.

Krumlauf R, Wilkinson DG. 2021. Segmentation and patterning of the vertebrate hindbrain. Development 148:dev186460.

Lancaster MA, Renner M, Martin CA, Wenzel D, Bicknell LS, Hurles ME, Homfray T, Penninger JM, Jackson AP, Knoblich JA. 2013. Cerebral organoids model human brain development and microcephaly. Nature 501: 373–379.

Laureano AS, Flaherty K, Hinman AM, Jadali A, Nakamura T, Higashijima SI, Sabaawy HE, Kwan KY. 2022. shox2 is required for vestibular statoacoustic neuron development. Biol Open 11:bio059599.

Li C, Xu D, Ye Q, Hong S, Jiang Y, Liu X, Zhang N, Shi L, Qin CF, Xu Z. 2016. Zika Virus Disrupts Neural Progenitor Development and Leads to Microcephaly in Mice. Cell Stem Cell 19: 672.

Li Q, Shen L. 2022. Neuron segmentation using 3D wavelet integrated encoder-decoder network. Bioinformatics 38: 809–817.

Lin W, Wang Y, Chen Y, Wang Q, Gu Z, Zhu Y. 2021a. Role of Calcium Signaling Pathway-Related Gene Regulatory Networks in Ischemic Stroke Based on Multiple WGCNA and Single-Cell Analysis. Oxid Med Cell Longev 2021: 8060477.

Lin WW, Xu LT, Chen YS, Go K, Sun C, Zhu YJ. 2021. Single-Cell Transcriptomics-Based Study of Transcriptional Regulatory Features in the Mouse Brain Vasculature. Biomed Res Int 2021: 7643209.

Lin YT, Seo J, Gao F, Feldman HM, Wen HL, Penney J, Cam HP, Gjoneska E, Raja WK, Cheng J, Rueda R, Kritskiy O, Abdurrob F, Peng Z, Milo B, Yu CJ, Elmsaouri S, Dey D, Ko T, Yankner BA, Tsai LH. 2018. APOE4 Causes Widespread Molecular and Cellular Alterations Associated with Alzheimer’s Disease Phenotypes in Human iPSC-Derived Brain Cell Types. Neuron 98: 1141–1154.e1147.

Lucke-Wold BP, Logsdon AF, Nguyen L, Eltanahay A, Turner RC, Bonasso P, Knotts C, Moeck A, Maroon JC, Bailes JE, Rosen CL. 2018. Supplements, nutrition, and alternative therapies for the treatment of traumatic brain injury. Nutr Neurosci 21: 79–91.

Madhavan M, Nevin ZS, Shick HE, Garrison E, Clarkson-Paredes C, Karl M, Clayton BLL, Factor DC, Allan KC, Barbar L, Jain T, Douvaras P, Fossati V, Miller RH, Tesar PJ. 2018. Induction of myelinating oligodendrocytes in human cortical spheroids. Nat Methods 15: 700–706.

Mattei PL, Beachkofsky TM, Gilson RT, Wisco OJ. 2012. Syphilis: a reemerging infection. Am Fam Physician 86: 433–440.

McNulty CL, Peres JN, Bardine N, van den Akker WM, Durston AJ. 2005. Knockdown of the complete Hox paralogous group 1 leads to dramatic hindbrain and neural crest defects. Development 132: 2861–2871.

Medoro AK, Sánchez PJ. 2021. Syphilis in Neonates and Infants. Clin Perinatol 48: 293–309.

Menon V, Thomas R, Elgueta C, Horl M, Osborn T, Hallett PJ, Bartos M, Isacson O, Pruszak J. 2019. Comprehensive Cell Surface Antigen Analysis Identifies Transferrin Receptor Protein-1 (CD71) as a Negative Selection Marker for Human Neuronal Cells. Stem Cells 37: 1293–1306.

Mercurio S, Serra L, Nicolis SK. 2019. More than just Stem Cells: Functional Roles of the Transcription Factor Sox2 in Differentiated Glia and Neurons. Int J Mol Sci 20:4540.

Mertens J, Marchetto MC, Bardy C, Gage FH. 2016. Evaluating cell reprogramming, differentiation and conversion technologies in neuroscience. Nat Rev Neurosci 17: 424–437.

Meyer U, Yee BK, Feldon J. 2007. The neurodevelopmental impact of prenatal infections at different times of pregnancy: the earlier the worse? Neuroscientist 13: 241–256.

Nelson SB, Hempel C, Sugino K. 2006. Probing the transcriptome of neuronal cell types. Curr Opin Neurobiol 16: 571–576.

Nowakowski TJ, Pollen AA, Di Lullo E, Sandoval-Espinosa C, Bershteyn M, Kriegstein AR. 2016. Expression Analysis Highlights AXL as a Candidate Zika Virus Entry Receptor in Neural Stem Cells. Cell Stem Cell 18: 591–596.

Pollen AA, Bhaduri A, Andrews MG, Nowakowski TJ, Meyerson OS, Mostajo-Radji MA, Di Lullo E, Alvarado B, Bedolli M, Dougherty ML, Fiddes IT, Kronenberg ZN, Shuga J, Leyrat AA, West JA, Bershteyn M, Lowe CB, Pavlovic BJ, Salama SR, Haussler D, Eichler EE, Kriegstein AR. 2019. Establishing Cerebral Organoids as Models of Human-Specific Brain Evolution. Cell 176: 743–756.e717.

Qian X, Nguyen HN, Song MM, Hadiono C, Ogden SC, Hammack C, Yao B, Hamersky GR, Jacob F, Zhong C, Yoon KJ, Jeang W, Lin L, Li Y, Thakor J, Berg DA, Zhang C, Kang E, Chickering M, Nauen D, Ho CY, Wen Z, Christian KM, Shi PY, Maher BJ, Wu H, Jin P, Tang H, Song H, Ming GL. 2016. Brain-Region-Specific Organoids Using Mini-bioreactors for Modeling ZIKV Exposure. Cell 165: 1238–1254.

Ramani A, Müller L, Ostermann PN, Gabriel E, Abida-Islam P, Müller-Schiffmann A, Mariappan A, Goureau O, Gruell H, Walker A, Andrée M, Hauka S, Houwaart T, Dilthey A, Wohlgemuth K, Omran H, Klein F, Wieczorek D, Adams O, Timm J, Korth C, Schaal H, Gopalakrishnan J. 2020. SARS-CoV-2 targets neurons of 3D human brain organoids. EMBO J 39: e106230.

Rubio-Hernández EI, Comas-García M, Coronado-Ipiña MA, Colunga-Saucedo M, González Sánchez HM, Castillo CG. 2023. Astrocytes derived from neural progenitor cells are susceptible to Zika virus infection. PLoS One 18: e0283429.

Saliba AE, Li L, Westermann AJ, Appenzeller S, Stapels DA, Schulte LN, Helaine S, Vogel J. 2016. Single-cell RNA-seq ties macrophage polarization to growth rate of intracellular Salmonella. Nat Microbiol 2: 16206.

Satija R, Farrell JA, Gennert D, Schier AF, Regev A. 2015. Spatial reconstruction of single-cell gene expression data. Nat Biotechnol 33: 495–502.

Setoh YX, Amarilla AA, Peng NYG, Griffiths RE, Carrera J, Freney ME, Nakayama E, Ogawa S, Watterson D, Modhiran N, Nanyonga FE, Torres FJ, Slonchak A, Periasamy P, Prow NA, Tang B, Harrison J, Hobson-Peters J, Cuddihy T, Cooper-White J, Hall RA, Young PR, Mackenzie JM, Wolvetang E, Bloom JD, Suhrbier A, Khromykh AA. 2019. Determinants of Zika virus host tropism uncovered by deep mutational scanning. Nat Microbiol 4: 876–887.

Sherman BT, Hao M, Qiu J, Jiao X, Baseler MW, Lane HC, Imamichi T, Chang W. 2022. DAVID: a web server for functional enrichment analysis and functional annotation of gene lists (2021 update). Nucleic Acids Res 50: W216–w221.

Shoja MM, Johal J, Oakes WJ, Tubbs RS. 2018. Embryology and pathophysiology of the Chiari I and II malformations: A comprehensive review. Clin Anat 31: 202–215.

Sugino K, Clark E, Schulmann A, Shima Y, Wang L, Hunt DL, Hooks BM, Tränkner D, Chandrashekar, J, Picard S, Lemire AL, Spruston N, Hantman AW, Nelson SB. 2019. Mapping the transcriptional diversity of genetically and anatomically defined cell populations in the mouse brain. Elife 8:e38619.

Tantalo LC, Lukehart SA, Marra CM. 2005. Treponema pallidum strain-specific differences in neuroinvasion and clinical phenotype in a rabbit model. J Infect Dis 191: 75–80.

Tong ML, Zhao Q, Liu LL, Zhu XZ, Gao K, Zhang HL, Lin LR, Niu JJ, Ji ZL, Yang TC. 2017. Whole genome sequence of the Treponema pallidum subsp. pallidum strain Amoy: An Asian isolate highly similar to SS14. PLoS One 12: e0182768.

Tonni G, Grisolia G, Pisello M, Zampriolo P, Fasolato V, Sindico P, Araújo Junior E, Bonasoni MP. 2022. Congenital Syphilis Presenting with Brain Abnormalities at Neuroscan: A Case Report and a Brief Literature Review. Microorganisms 10:1497.

Trujillo CA, Muotri AR. 2018. Brain Organoids and the Study of Neurodevelopment. Trends Mol Med 24: 982–990.

Valakh V, Wise D, Zhu XA, Sha M, Fok J, Van Hooser SD, Schectman R, Cepeda I, Kirk R, O’Toole SM, Nelson SB. 2023. A transcriptional constraint mechanism limits the homeostatic response to activity deprivation in mammalian neocortex. Elife 12: e74899.

Velasco S, Kedaigle AJ, Simmons SK, Nash A, Rocha M, Quadrato G, Paulsen B, Nguyen L, Adiconis X, Regev A, Levin JZ, Arlotta P. 2019. Individual brain organoids reproducibly form cell diversity of the human cerebral cortex. Nature 570: 523–527.

Verghese VP, Hendson L, Singh A, Guenette T, Gratrix J, Robinson JL. 2018. Early Childhood Neurodevelopmental Outcomes in Infants Exposed to Infectious Syphilis In Utero. Pediatr Infect Dis J 37: 576–579.

Watanabe M, Buth JE, Vishlaghi N, de la Torre-Ubieta L, Taxidis J, Khakh BS, Coppola G, Pearson CA, Yamauchi K, Gong D, Dai X, Damoiseaux R, Aliyari R, Liebscher S, Schenke-Layland K, Caneda C, Huang EJ, Zhang Y, Cheng G, Geschwind DH, Golshani P, Sun R, Novitch BG. 2017. Self-Organized Cerebral Organoids with Human-Specific Features Predict Effective Drugs to Combat Zika Virus Infection. Cell Rep 21: 517–532.

Wicher V, Wicher K. 2001. Pathogenesis of maternal-fetal syphilis revisited. Clin Infect Dis 33: 354–363.

Woods CR. 2009. Congenital syphilis-persisting pestilence. Pediatr Infect Dis J 28, 536–537.

Yang SG, Wang XW, Qian C, Zhou FQ. 2022. Reprogramming neurons for regeneration: The fountain of youth. Prog Neurobiol 214: 102284.

Zhang M, Ge T, Zhang Y, La X. 2023. Identification of MARK2, CCDC71, GATA2, and KLRC3 as candidate diagnostic genes and potential therapeutic targets for repeated implantation failure with antiphospholipid syndrome by integrated bioinformatics analysis and machine learning. Front Immunol 14: 1126103.

Zhang T, Lin Y, Liu J, Zhang ZG, Fu W, Guo LY, Pan L, Kong X, Zhang MK, Lu YH, Huang ZR, Xie Q, Li WH, Xu XQ. 2016. Rbm24 Regulates Alternative Splicing Switch in Embryonic Stem Cell Cardiac Lineage Differentiation. Stem Cells 34: 1776–1789.

Zhang W, Tan YW, Yam WK, Tu H, Qiu L, Tan EK, Chu JJH, Zeng L. 2019. In utero infection of Zika virus leads to abnormal central nervous system development in mice. Sci Rep 9: 7298.

