## Supplemental File for "Single-cell RNA sequencing of iPSC-derived brain organoids reveals *Treponema pallidum* infection inhibiting neurodevelopment"

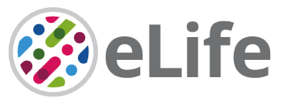

Supplementary File for

**Single-cell RNA sequencing of iPSC-derived brain organoids reveals *Treponema pallidum* infection inhibiting neurodevelopment**

**This Supplement file includes:**

Abbreviations

Figure supplememt 1 to 4

**Abbreviations:**

Ve-cad: Vascular endothelial cadherin; KDR: Kinase insert domain receptor; SOX17: Sex determining region Y-box 17; MAP2: Microtubule associated protein 2; SOX2: Sex determining region Y-box 2; TUBB3: Tubulin beta 3 class III; HOXB2: Homeobox B2; HOXA2: Homeobox A2; HOTAIRM1: HOX antisense intergenic RNA myeloid 1; IGFBP1: Insulin like growth factor binding protein 1; EN2: Engrailed homeobox 2; SPRY1: Sprouty 1; HOXC4: Homeobox C4; HOXC5: Homeobox C5; HOXA5: Homeobox A5; NEUROD: Neurogenic differentiation 4; NEUROG1: Neurogenin 1; PCP4: Purkinje cell protein 4; GNG5: G protein subunit gamma 5; HOXA4: HOXA5: Homeobox A4; MAGEH1: Melanoma antigen family H 1; MEIS3: Meis homeobox 3; USP47: Ubiquitin specific peptidase 47; SHOX2: Short stature homeobox 2; HES4: Hairy and enhancer of split 4; ASPM: Assembly factor for spindle microtubules; CCNB1: CyclinB1; TPX2: Targeting protein for xenopus kinesin-like protein2; UBE2C: Ubiquitin conjugating enzyme E2 C; TOP2A: Topoisomerase II alpha; MKI67: antigen identified by monoclonal antibody Ki-67; CENPF: Centromere protein F; PTTG1: Pituitary tumor-transforming 1; HMGN2: High mobility group nucleosomal binding domain 2; SNCG: Synuclein gamma; S100A10: S100 calcium binding protein A10; RGS10: Regulator of G protein signaling 10; STMN2: Stathmin-like 2; PPP1R17: Protein phosphatase 1 regulatory subunit 17; BASP1: Brain abundant membrane attached signal protein 1; NEFL: Neurofilament light chain; CALM1: Calmodulin 1; DCX: Doublecortin; TMSB10: Thymosin beta 10; TMSB4X: Thymosin beta 4 X-linked; STMN4: Stathmin-like 4; C11orf96: Chromosome 11 open reading frame 96; TUBA1A: tubulin alpha 1a; RTN1: Reticulon 1; SYT4: Synaptotagmin 4; SST: Somatostatin; LRRC75A-AS: Leucine-rich repeat-containing protein75A-antisense1; ZFAS1: Zinc finger protein antisense strand 1; GAS5: Growth arrest specific 5; ID1: Inhibitor of DNA binding 1; NDUFB5: NADH dehydrogenase (ubiquinone) 1 beta subcomplex 5; BNIP3: BCL2 interacting protein 3; PGK1: Phosphoglycerate Kinase 1; IFITM3: Interferon induced transmembrane protein 3; POU3F3: POU class 3 homeobox 3; INSIG2: Insulin induced gene 2; NEFM: Neurofilament medium chain; NEFL: Neurofilament light chain; ENO2: Enolase 2; B4ALT5: Beta-1,4-galactosyltransferase 5; SPTAN1: Spectrin alpha, non-erythrocytic 1; TTR: Transthyretin.

**Figure supplement 1: Construction and verification of a three-dimensional culture system for brain organoids.**

In order to better imitate the process of brain nerve development, a three-dimensional culture system of brain organoids was constructed. iPSC with good growth status was centrifuged to obtain precipitated cell spheres, gently resuspended in brain organoid (Stage I) medium, and then placed in a low-adsorption culture plate for suspension culture for 3 days to form multiple miniature neurosphere aggregates, namely embryoid bodies. Continued culture to day 7, the volume of the embryoid body gradually increased and formed a sac cavity, which looked like an organoid under the microscope. At this time, the neurosphere gradually formed neuroectoderm, the neural differentiation was started immediately, and then change culture medium (stage II), which was conducive to the differentiation and development of the nerve. At this stage, the volume of the organoid gradually increased and the shape of the sac gradually became clear with the passage of time. The early nerve rosette structure could be observed on day 30. Then, the culture medium (Stage III) was changed, and relatively complete and mature neural roseate structure appeared after day 45 (Figure supplement 1A).

In order to verify the construction process of brain organoids, we collected the cells (or microspheres, organoids) of day 0, day 7, day 14, day 30 and day 45 at 5 time points during the formation of organoids, and detected the transcriptome levels. The results showed that the mRNA expression of nestin, an early marker of neural progenitor cells, increased slowly at the beginning, reached the peak on the 14th day, and then gradually decreased. PAX6, a late marker of neural progenitor cells, began to express on day 7, gradually increased with the passage of culture time until day 45, and then remained a plateau. The neuron-specific cytoskeletal protein MAP2 was weakly expressed from day 14 and rapidly increased on day 30, indicating that the late organoids had gradually differentiated and matured (Figure supplement 1B). Furthermore, immunofluorescence staining on frozen sections of organoids showed that SOX2 was scattered on day 30, but several relatively complete structures of neural rosettes were observed on day 45 (Figure supplement 1C). The above results show that we have successfully constructed brain organoids, which can be used for subsequent experimental research.

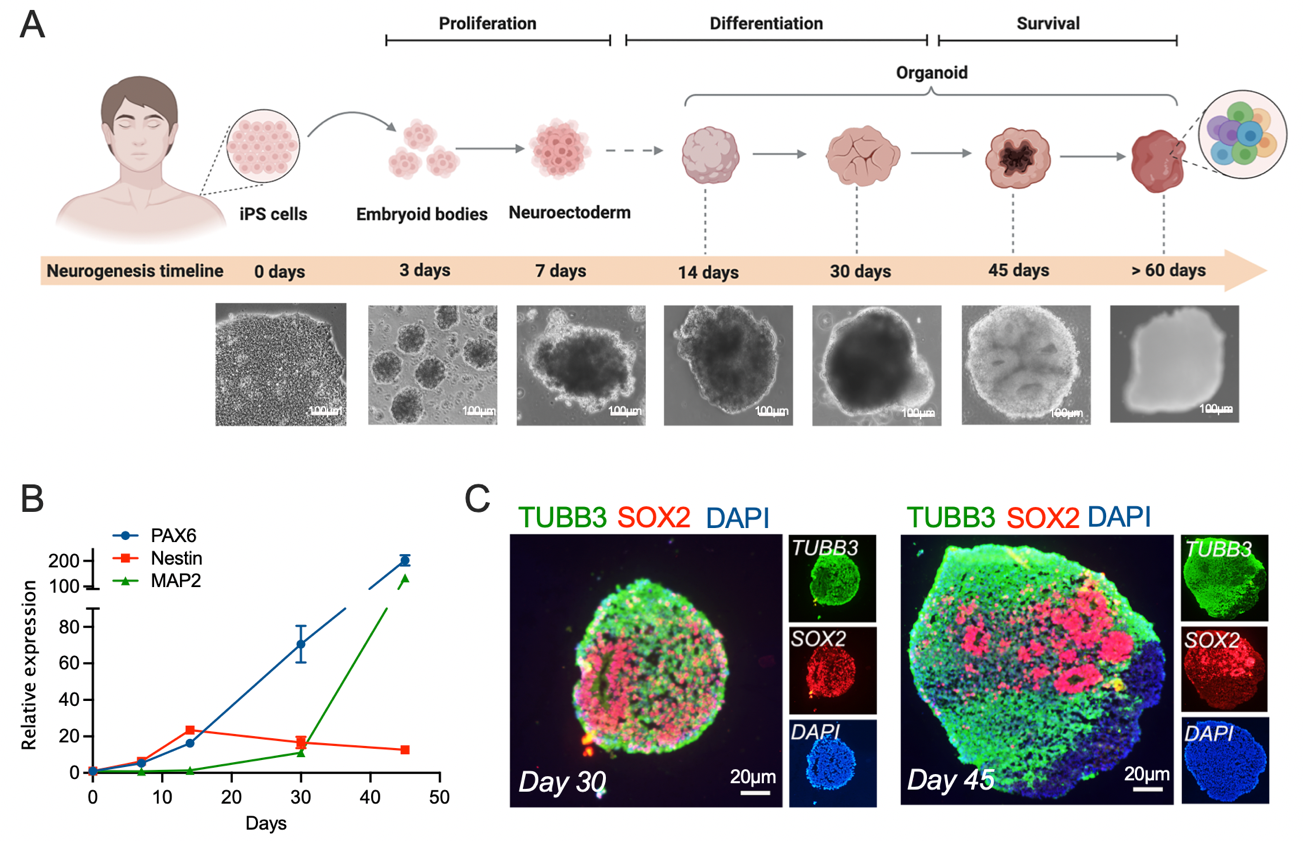

**Figure supplement 1.** Construction and verification of a three-dimensional culture system for brain organoids. (**A**) Schematic diagram of brain organoid three-dimensional culture system. (**B**) mRNA expression changes of neural markers during the construction of three-dimensional culture system. (**C**) Immunofluorescence of organoids frozen sections.

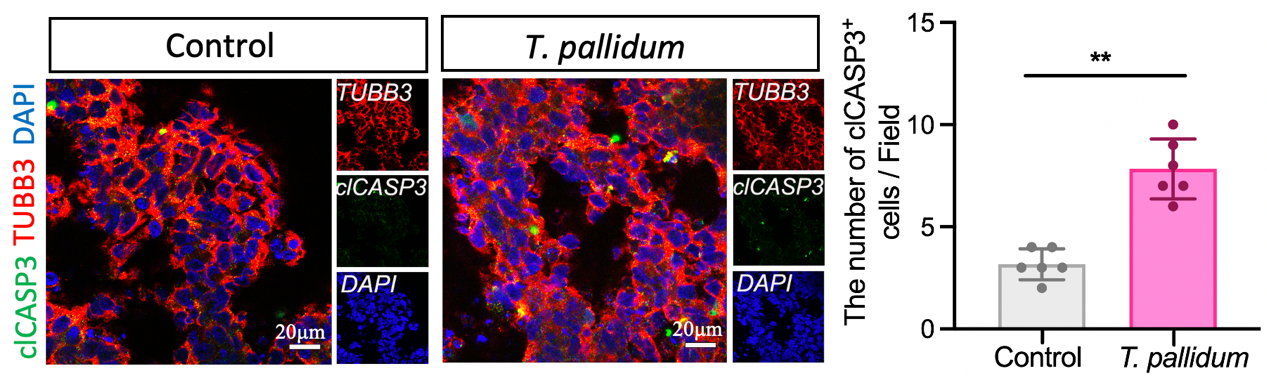

**Figure supplement 2.** The number of clCASP3^+^ cells in the microscopic field of brain organoids. A nonparametric *t*-test was used to evaluate the statistical differences between the two groups. (**: *P* < 0.01).

**Figure supplement 3: Definition, naming and evaluation of each cell population based on scRNA-seq**

Representative marker genes with the highest expression in each cell population were selected to define each cell population based on scRNA-seq. The results showed that marker genes with specific expression exist in all other groups except for the fifth group (Figure supplement 2A), indicating the reliability of scRNA-seq data. Furthermore, two marker genes were selected from the cell types of each neurodevelopmental stage to define each cell population according to the published database (based on the control group data) (Figure supplement 2B). *DCT* and *HMX1* marker genes were used to define the melanocytic cell population, *HOXD4* and *PCDH8* marker genes were used to define the neural progenitor cell population, *NRXN1* and *SLC17A6* marker genes were used to define neurones, and *EMX1* and *FABP7* marker genes were used to define glia cells. *CBLN1* and *FEZF1* marker genes were used to define neural ectodermal cells. *ANKRD37* and *STC1* marker genes were used to define neural epithelial cells. *UBE2C* and *TK1* marker genes were used to define neuronal stem cells. *PMCH* and *HTR2C* marker genes were used to define cilia like cells. *CYP1A1* and *AP1M2* marker genes were used to define ectodermal cells. *SOX10* and *GJC3* marker genes were used to define oligodendrocytes. *COL6A3* and *LUM* marker genes were used to define mesenchymal stem cells. A total of 12 cell groups were named and defined, and neural progenitor cells were divided into neural progenitor cell subgroup 1 and subgroup 2, and one cell group was not named and defined.

In order to verify the specificity of cell group defined above, we performed cluster correlation analysis on the specific genes of each cell group, which was reflected in the form of heatmap. The positive and negative genes were represented using yellow and purple colours, respectively. The heatmap analysis was performed on five signature genes for each type of nerve cell. The results demonstrated a high correlation between the signature genes and cell types defined by each population (Figure supplement 2C). The above results indicate that each cell population that was defined and named according to scRNA-seq data is reasonably matched and could be used for subsequent analysis and research.

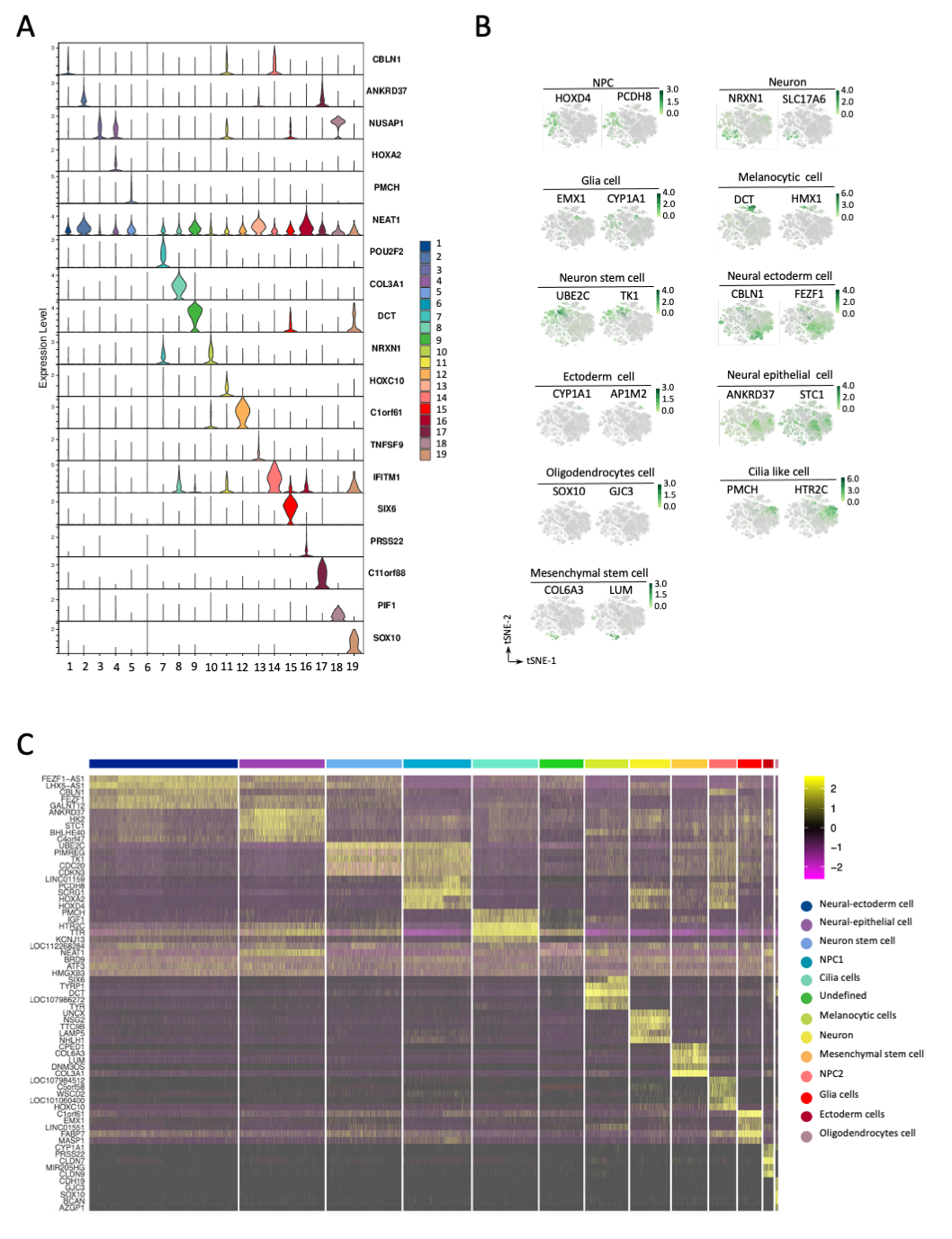

**Figure supplement** **3.** Definition, naming and evaluation of each cell population according to different marker genes. (**A**) Violin plot of different marker genes for each cell population. (**B**) tSNE distribution map of each cell population calibrated by published marker genes. (**C**) Heat map specificity analysis of each cell population.

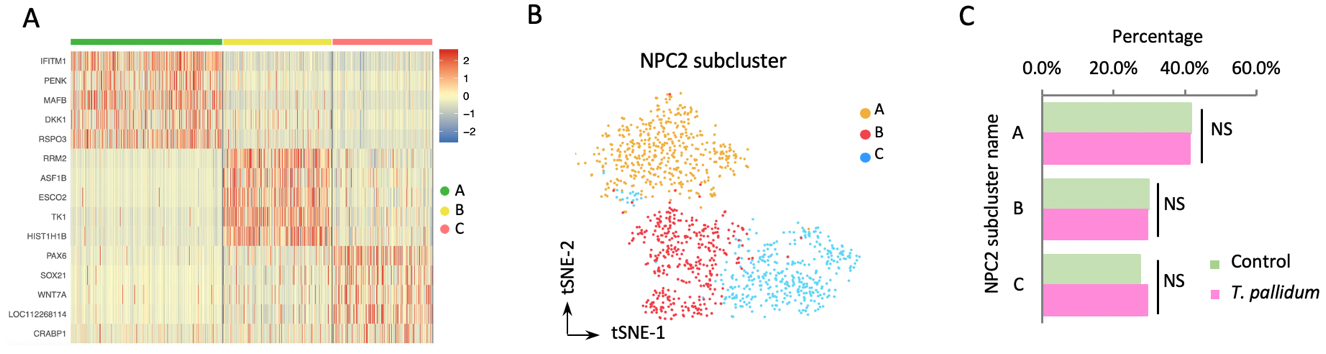

**Figure supplement 4.** *T. pallidum* did not affect the differentiation of neural progenitor cell subset 2 in brain organoids. (**A**) The correlation and expression differences of marker genes in NPC2 groups were analyzed by heat map. (**B**) tSNE dimensional reduction location distribution map of NPC2 group. (**C**) The proportion changes of the three small clusters before and after *T. pallidum* infection. The Chi-square test was used to compare the percentage differences among different groups (NS: not significant).
